# Monocyte Metabolic Plasticity and Cytokine Production Differentiate Latent TB Infection from Active Disease

**DOI:** 10.1101/2025.04.24.650431

**Authors:** Gráinne Jameson, Isabella Batten, Adam H. Dyer, Caoimhe Geoghegan, Moninne Murray, Niamh McDonnell, Dearbhla M. Murphy, Sarah A. Connolly, Anne Marie McLaughlin, Cilian Ó Maoldomhnaigh, Laura E. Gleeson, Joseph Keane, Sharee A. Basdeo

**Affiliations:** School of Medicine, Trinity Translational Medicine Institute, St James’s Hospital, Trinity College Dublin, Dublin, Ireland; National Tuberculosis Centre, Department of Respiratory Medicine, St James’s Hospital, Dublin, Ireland; Respiratory Department, St James’s Hospital, Dublin 8, Dublin, Ireland

**Keywords:** Mycobacterium tuberculosis, Mitochondria, Innate Immunity, monocyte

## Abstract

**Rationale:** Monocytes are central to host defence against *Mycobacterium tuberculosis* (Mtb), yet their functional and metabolic profiles during latent TB infection (TBI) and active TB disease (TBD) remain poorly defined. Immunometabolic dysfunction may underlie ineffective responses in TB, but cell-specific mechanisms are unclear.

**Objectives:** To compare the phenotypic, functional, and metabolic profiles of circulating monocytes from individuals with TBI, TBD, and healthy controls (HC), and assess the impact of treatment.

**Measurements:** Peripheral blood monocytes were profiled using high-dimensional flow cytometry, Luminex cytokine/chemokine assays, and SCENITH™, a flow-based metabolic assay. Unstimulated and Mtb-stimulated monocytes from treatment-naïve and treated individuals were analysed.

**Main Results:** Monocytes from TBI and TBD showed distinct phenotypes from HC, marked by elevated CD14 and CD45RA. HLA-DR was reduced in TBI versus HC and further decreased in TBD. TNF receptors were downregulated in TBI but unchanged in TBD. Baseline cytokine and chemokine profiles in TBI and TBD were similar (yet distinct from HC), but Mtb stimulation elicited a stronger cytokine response in TBI. Metabolically, TBI and TBD monocytes exhibited increased glycolysis and reduced mitochondrial dependence versus HC. Treatment partially restored mitochondrial function. Upon Mtb challenge, TBI monocytes had higher glycolytic capacity than TBD.

**Conclusions:** Monocyte metabolic plasticity and cytokine production distinguish latent from active TB and are partially reversible with treatment. Circulating monocyte metabolism reflects TB immune status and may serve as a biomarker or therapeutic target. Reprogrammed glycolytic profiles in TBI contrast with impaired adaptability in TBD, suggesting dysfunctional myeloid activation during disease.

## Introduction

Tuberculosis (TB) is the leading cause of death by an infectious agent, with an estimated 1.25 million deaths and 10.8 million cases in 2023 (1). While most individuals infected with *Mycobacterium tuberculosis* (Mtb) do not progress to active disease, those with latent TB infection (TBI) carry approximately twice the risk of developing tuberculosis disease (TBD compared with uninfected individuals), yet also demonstrate a 79% lower risk of progressive disease following reinfection (2). Understanding the immunological mechanisms that confer protection in TBI or promote progression to TBD is a central question in TB research.

Monocytes are short-lived, circulating myeloid cells that play a critical role in the peripheral immune response to infection. They contribute to antimicrobial defence through phagocytosis, cytokine production, and antigen presentation (3–7). In TB, monocytes are recruited from the circulation to the lungs, the primary site of infection, where they can differentiate into alveolar macrophages (AMs) (8–12). The AM niche almost exclusively harbours early Mtb infection and acts as the first line of defence (13,14).

Cellular metabolism has emerged as a key regulator of myeloid function. Glycolysis, in particular, drives effective antimicrobial activity in both human AM and monocyte-derived macrophages responding to Mtb (15–18). Similar metabolic reprogramming occurs in healthy monocytes upon infection (19). In contrast, monocytes from individuals with TBD exhibit impaired ex vivo metabolic function, characterised by reduced glycolysis and oxidative phosphorylation compared with healthy controls (20). These deficits are partially reversible with TB treatment, suggesting that Mtb infection directly alters monocyte bioenergetics and function.

Together, these findings highlight the importance of immunometabolism in shaping host responses to TB and support ongoing efforts to develop host-directed therapies. However, how monocyte metabolism and function differ between stages of TB, particularly between TBI and TBD, remains incompletely understood. In this study, we aimed to characterise the phenotypic, functional, and metabolic profiles of circulating monocytes from individuals across the TB spectrum. We used SCENITH™, a single-cell metabolic profiling assay, to uncover immune and bioenergetic features that may distinguish infection from disease(21) (Figure 1).

**Figure 1.**
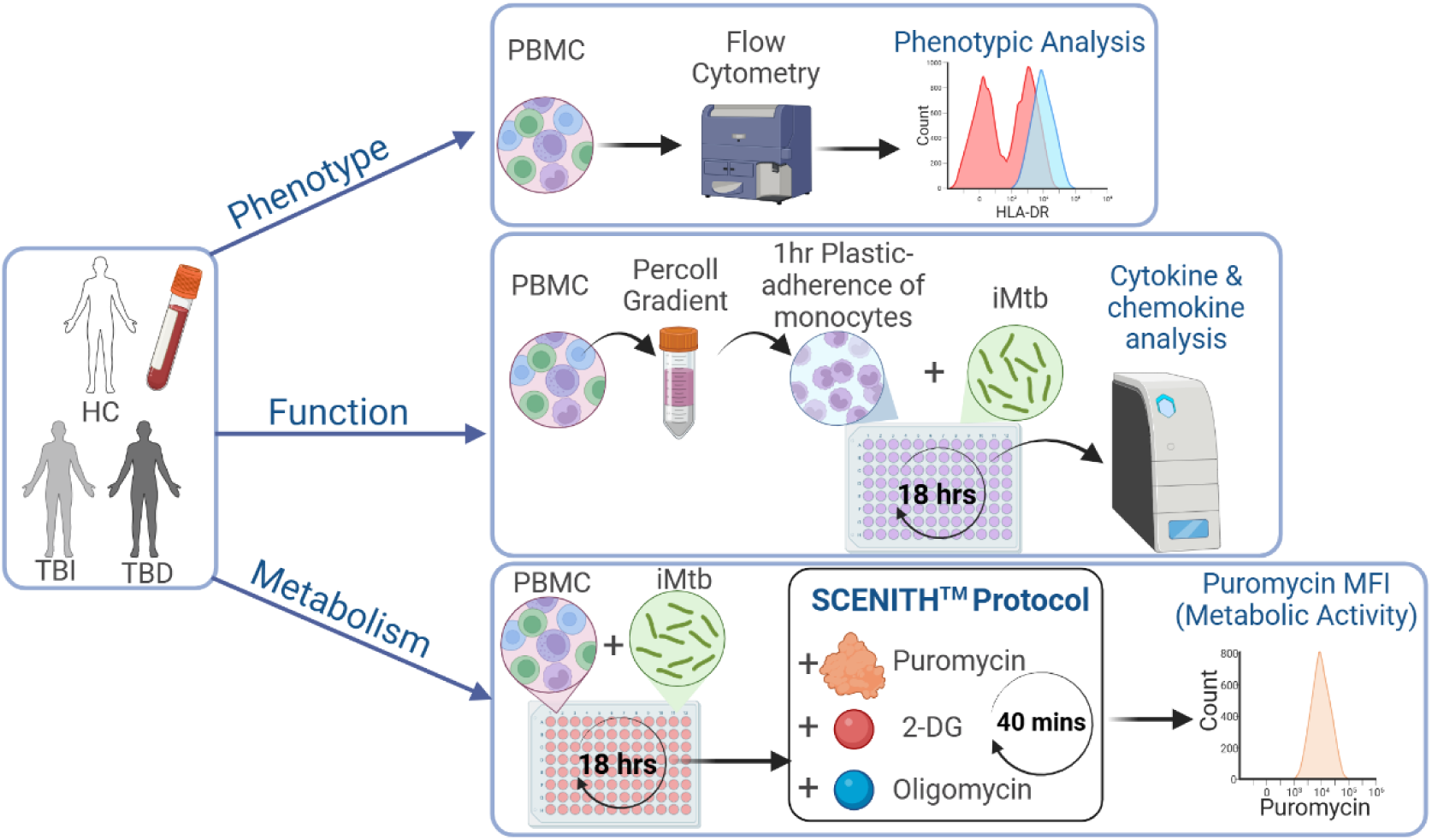
Schematic of study design for analysis of monocyte phenotype, function and metabolism. Peripheral blood samples were collected from healthy controls (HC), people with TBI and people with TBD and PBMC were isolated by density centrifugation. Isolated PBMCs were stained for flow cytometry analysis of monocyte expression levels of CD14, CD45RA, CD40, HLA-DR, TNFRI and TNFRII. Monocytes were isolated from PBMCs using density centrifugation with a Percoll gradient followed by 1hr plastic adherence. Purified monocytes were then cultured, with or without irradiated-H37Rv (iMtb), for 18 hours. The supernatants were harvested and analysed for cytokine and chemokine concentrations. PBMC were plated for 24 hours with or without iMtb. Puromycin and metabolic inhibitors (2-DG and oligomycin) were added for 40 mins and then the cells were stained for flow cytometry analysis. Cells were washed and stained with fluorochrome-conjugated antibodies against CD14 and puromycin and analyzed by flow cytometry. Puromycin MFI levels measure overall protein synthesis and act as a surrogate for metabolic activity. Created in https://BioRender.com.

## Methods

### Human Participants

All peripheral blood patient samples were collected in the out-patient department of St James’s Hospital Dublin when patients were attending routine appointments (Table 1). Written informed consent was obtained prior to collection. Exclusion criteria included age under 18 years, inability to provide written informed consent, multi-drug-resistant tuberculosis, or a known (or ensuing) diagnosis of malignancy, HIV or any other infection. Individuals with TBI were selected based on a positive QuantiFERON-TB Gold Plus IGRA test with absence of any clinical symptoms of TB disease and were subdivided into those pre-treatment (TBI/-; n=6) and those on-treatment (TBI/+; n=4). Individuals clinically diagnosed of TBD were grouped into intensive (TBD/I, <2 months treatment of 3-4 drugs; n=4) and continuation (TBD/C, >2 months reduced to 1-2 drugs; n=4) treatment phases. Peripheral blood samples were also obtained from healthy controls who have all received BCG vaccination previously.

**Table 1:** Patient information. HC=Healthy Controls, TBI= Tuberculosis Infection, TBD= tuberculosis disease, F= female, M=male.

|  | HC | TBI | TBD |
| --- | --- | --- | --- |
| **Age** | 38+/-15 | 44.90+/-12.81 | 41.25+/-18.43 |
| **Gender** | F (9) M (1) | M (11) F (10) | M (10) F (6) |
| **Ethnicity**<br>**(self-reported)** | European (9) African (1) | Asian (10), European (4), African (2), Indian (2), Not reported (3) | Asian (4), European (5), African (7), |
| **Site of Mtb infection** | N/A | N/A | Pulmonary (6), Extra-pulmonary (7), Both (2) |

### PBMC and Monocyte Isolations

PBMC were isolated from peripheral blood samples by density centrifugation over Lymphoprep (StemCell Technologies). Monocytes were separated by hyperosmotic density gradient centrifugation by layering PBMC on top of a fresh hyperosmotic Percoll solution (48.5% Percoll, 41.5% sterile H2O, 0.16 M NaCl) and centrifuged for 15 minutes at 580*g*. The interphase layer was isolated, and cells were washed with cold PBS and resuspended at a concentration of 1 × 10^6^ cells/mL. Cells were plastic adhered for 1hr in serum-free medium and washed twice with PBS to remove any non-adherent cells.

Irradiated H37Rv (iH37Rv; gifted by BEI Resources) stocks were thawed, sonicated for 15 mins and syringed three times through a 25G needle break up clumps. iH37Rv was centrifuged at 30 xg for 5 mins to pellet any further clumps. Supernatant was aspirated and the concentration of iH37Rv was calculated using Pierce^TM^ BCA Protein Assay with Albumin protein as a control (Thermo Scientific^TM^). Cells were stimulated with 10 µg/mL of iH37Rv overnight for 18hours. Cells were left unstimulated as a control.

### Cytokine and Chemokine Analyses

A Human Luminex Discovery Kit (Cat LXSAHM-25) was designed to assess the following analytes in monocyte supernatants: CCL1/I-309/TCA-3 (BR15), CCL2/JE/MCP-1 (BR25), CCL3/MIP-1 alpha (BR35), CCL4/MIP-1 beta (BR37), CXCL1/GRO alpha/KC/CINC-1 (BR77), CXCL2/GRO beta/MIP-2/CINC-3 (BR27), CXCL9/MIG (BR52), CXCL10/IP-10/CRG-2 (BR67), G-CSF (BR54), GM-CSF (BR46), Granzyme B (BR57), IFN-alpha (BR63), IFN-beta (BR21), IFN-gamma (BR29), IL-1 alpha/IL-1F1 (BR38), IL-1 beta/IL-1F2 (BR28), IL-1ra/IL-1F3 (BR30), IL-6 (BR13), IL-8/CXCL8 (BR18), IL-10 (BR22), IL-12 p70 (BR56), IL-17/IL-17A (BR42), IL-18/IL-1F4 (BR78), IL-23 (BR76), TNF-alpha (BR12). The assay was carried out as per manufacturers protocol and analysed on Luminex MAGPIX System. Unstimulated and Mtb-stimulated cells from each donor were analyzed concurrently on the same plate, thereby eliminating potential inter-assay variability and ensuring the accurate calculation of fold-change values. Analyte concentrations were determined based on standards supplied in the kit, with a 5 parameter logistic regression (5-PL) curve used on xPONENT(TM) software.

### Flow cytometry

For phenotypical analyses PBMC were resuspended in PBS (Gibco) and stained immediately. Anti-CD86 BV421 (1:100, IT2.2), anti-CD45RA BV711 (1:100, HI100), anti-CD40 APCCy7 (1:100, 5C3), anti-HLADR APC (1:100, L243), anti-CD14 AlexaFlour 488 (1:100, HCD14), anti-CD16 PerCpCy5.5 (1:100, 3G8), anti-CD80 PeCy7 (1:100, 2D10), anti-TNFRI PE (1:50, W15099A), and anti-TNFRII PeDazzle594 (1:100, 3G7A02) were purchased from BioLegend and added to the cells for 10 mins at RT. Cells were washed, fixed and acquired or stored at 4°C until acquisition.

Cultured PBMC were resuspended in 100 μl of FcR block (1μl/test; Miltenyi Biotec) and ZombieNIR stain (BioLegend) in 1X PBS (Gibco). Cells were then surface stained with anti-CD14 and permeabilized with Foxp3 staining kit (eBioscience) according to the manufacturer’s instructions. Cells were incubated for 30 mins at 4°C with anti-puromycin (1:100, 2A4; BioLegend). Cells were acquired on a FACS Canto (BD Biosciences), BD Fortessa, (BD Biosciences) or an Amnis CellStream (Luminex Corporation, Austin, TX, USA). All data analysis was performed using FlowJo software (BD Biosciences).

### SCENITH (Single Cell mEtabolism by profiling Translation inHibition)

This protocol uses puromycin as a surrogate for protein synthesis, as it becomes incorporated into newly formed proteins. Since protein synthesis represents around 50% of total cellular metabolic activity, measuring puromycin incorporation provides a robust indicator of overall metabolic activity. By measuring puromycin incorporation post-oligomycin treatment we can estimate the cells dependency on mitochondrial respiration and their capacity for glycolysis. By measuring puromycin incorporation post 2-Deoxy-D-Glucose (2-DG) treatment we can estimate the cells dependency on glucose as a fuel and therefore their capacity for other fuels such as fatty acid and amino acid oxidation. A detailed explanation of these calculations was described in our previous work and in the original protocol publication (21,22). Post 18 hours of culture, SCENITH metabolic function profiling was carried out on PBMCs. Cells were treated for 40 mins at 37°C, 5% CO2 with DMSO Control (Co, 0.02% DMSO), 2-DG (100 mM; Sigma-Aldrich), Oligomycin (O; 1μM; Sigma-Aldrich), a combination of both drugs (DGO), or with Harringtonine, which inhibits protein synthesis (Sigma-Aldrich; 2 µg/ml). Puromycin (Puro; 10 μg/mL; Sigma-Aldrich) was added for the final 35 mins. Cells were washed in PBS and flow cytometry staining was carried out. The following calculations were used:

Co = GeoMFI of anti-Puro-Fluorochrome upon DMSO Control treatment.

DG = GeoMFI of anti-Puro-Fluorochrome upon 2-Deoxy-D-Glucose treatment.

O = GeoMFI of anti-Puro-Fluorochrome upon Oligomycin A treatment.

DGO = GeoMFI of anti-Puro-Fluorochrome upon DG+O treatment.

Mitochondrial Dependence (MD)=100(Co-O)/(Co-DGO)

Glycolytic Capacity (GC)= 100-100(Co-O)/(Co-DGO)

Fatty Acid and Amino Acid Oxidation Capacity (FAOC)= 100-(100(Co-DG)/(Co-DGO)

Glucose Dependence (GD)= 100(Co-DG)/(Co-DGO)

Therefore, GC is the inverse data of MD (i.e. GC= 100-MD) and FAOC is the inverse data of GD (i.e. FAOC= 100-GD). Patient samples with sufficient cell numbers underwent metabolic profiling. No data was omitted.

### Statistics

Statistical analysis was performed using GraphPad Prism version 10. The statistical test used is indicated in each figure legend. A P-value of <0.05 was considered significant.

### Study Approval

Our study protocol was approved by the Joint Research Ethics Committee of St. James’s and Tallaght Hospitals and by Trinity College Dublin. Written informed consent was received from participants prior to inclusion in the study.

### Data availability

Data is available directly from the corresponding author.

## Results

### Altered Expression of HLA-DR, CD14, and TNFRs in Monocytes Across the TB Spectrum

We next investigated whether TB infection influences monocyte frequency and phenotype across the disease spectrum. No significant differences were observed in the frequencies of classical, intermediate, or non-classical monocyte subsets between groups (Figure 2A–B). However, phenotypic analysis revealed elevated expression intensity of CD14 and CD45RA on classical monocytes from individuals with both TBI (CD14: P = 0.0039; CD45RA: P = 0.0070) and TBD (CD14: P = 0.0010; CD45RA: P = 0.0007) compared with healthy controls (HC) (Figure 2C–D).

**Figure 2.**
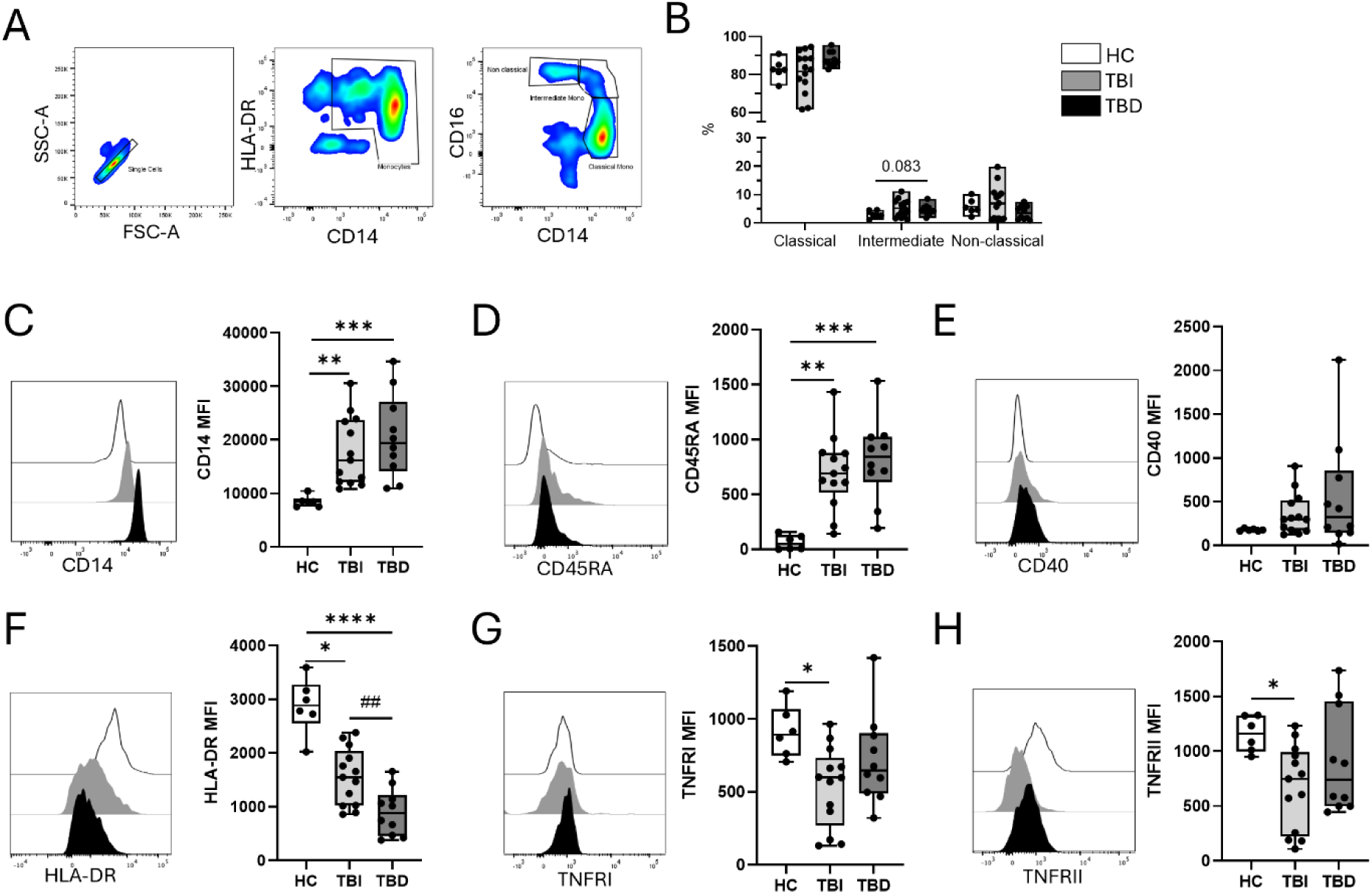
Distinct phenotype in monocytes from individuals with TBI. PBMC were isolated from peripheral blood and stained for flow cytometry analysis. **A-B)** Representative flow plot of gating strategy (A) and bar chart of monocyte subset frequencies as percentage of total monocytes (B). **C-H)** Expression levels of CD14 (C), CD45RA (D) and CD40 (E) and HLA-DR (F), TNFRI (G) and TNFRII (H) on classical monocytes across HC, TBI and TBD. P values generated using Two-Way ANOVA with Tukey’s multiple comparison test (B) and Kruskal Wallis test with Dunn’s multiple comparison (C-H), *P<0.05, **P<0.01, ***P<0.001, ****P<0.0001 or Mann Whitney test, ##, P<0.01 (F).

In contrast, expression of HLA-DR was significantly reduced on classical monocytes from TBI (P = 0.0374) and even further reduced in TBD (P < 0.0001) compared with HC, with significantly lower levels in TBD than TBI (P = 0.0068; Figure 2F). Expression of TNF receptors was decreased in TBI but not TBD; both TNFRI (P = 0.0185) and TNFRII (P = 0.0414) were downregulated on classical monocytes from TBI compared with HC (Figure 2G–H). Similar phenotypic changes were observed in intermediate and non-classical monocyte subsets across the TB spectrum (Supplemental Figure 2).

These data demonstrate that while monocyte subset frequencies remain stable, TB infection— both latent and active—induces distinct phenotypic reprogramming of monocytes. Notably, the progressive reduction in HLA-DR expression from TBI to TBD may reflect functional impairment, with implications for antigen presentation and immune regulation during infection (23–27).

### Monocytes from TBI and TBD individuals exhibit distinct functional responses to Mtb stimulation

To assess functional differences in monocytes across the TB spectrum, CD14^+^ monocytes were isolated from PBMC and cultured for 18 hours with or without irradiated-H37Rv (iMtb). Culture supernatants were collected, and cytokine and chemokine concentrations were measured using a custom Luminex multiplex bead-based immunoassay (Figure 3).

**Figure 3.**
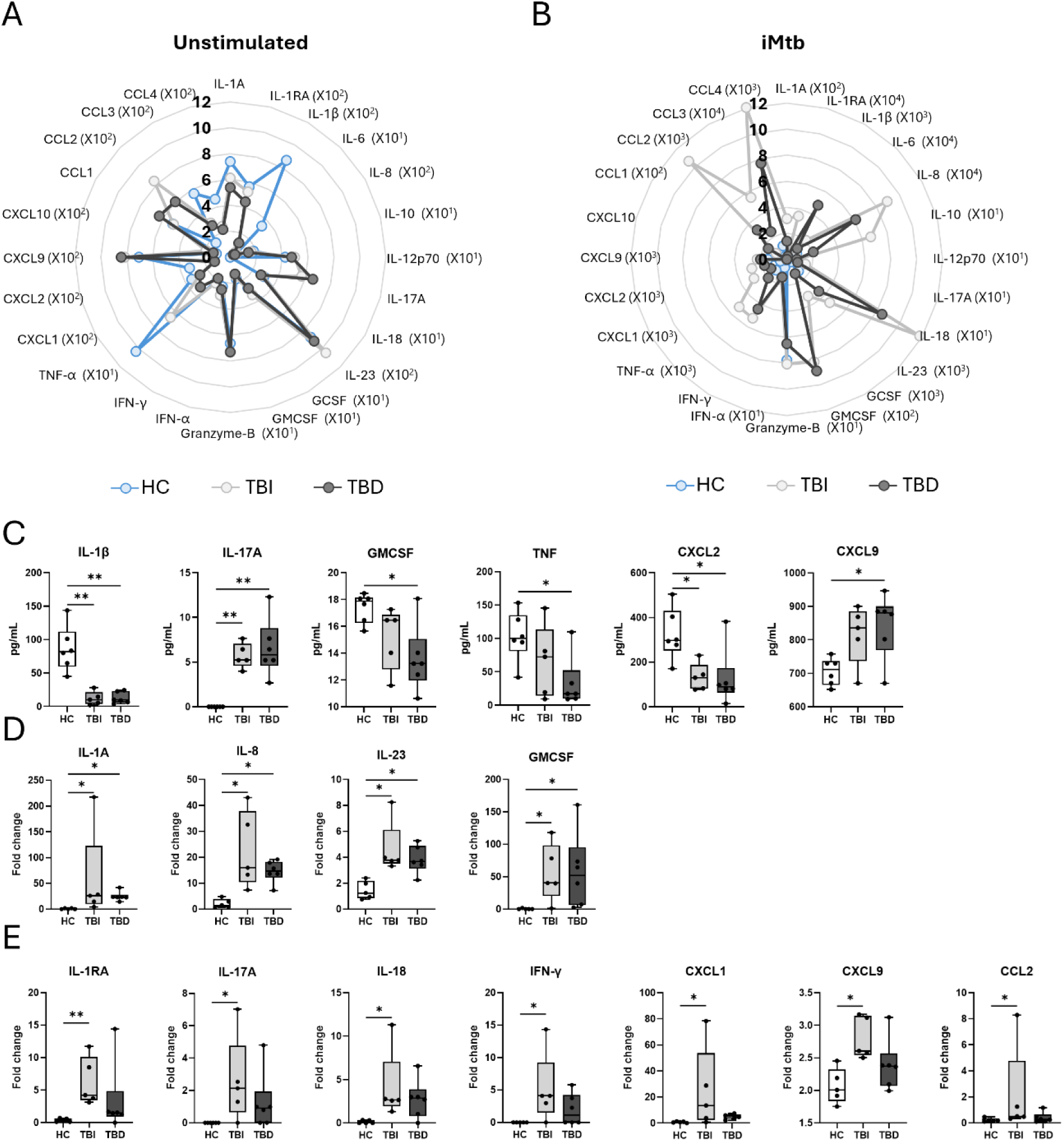
Monocytes from TBI individuals’ distinct cytokine and chemokine production in response to Mtb. Monocytes were isolated from PBMC and cultured for 18 hours with and without irradiated H37Rv (iMtb). [Healthy controls (HC, n=5)), latent (TBI, n=8) and Active (TBD, n=5) TB patients]. **A-B)** Mean levels of cytokine and chemokine concentrations (pg/mL) in supernatants collected from unstimulated (A) and iH37Rv (iMtb) stimulated (B) monocyte cultures [HC (blue), TBI (light grey) and TBD (dark grey]. **C)** Differential concentration of cytokines and chemokines in unstimulated monocytes. **D-E)** Analytes that are significantly increased in both TBI and TBD individual’s monocytes (D) and in only TBI individual’s monocytes (E). P values calculated using Kruskal Wallis test and Dunn’s multiple comparison test.

Unstimulated monocytes from individuals with latent TB infection (TBI, light grey) and active TB disease (TBD, dark grey) displayed cytokine/chemokine profiles that were distinct from healthy controls (HC, blue) (Figure 3A, C and Supplemental Figure 1A). Compared with TBD monocytes, unstimulated monocytes from HC produced significantly higher levels of IL-1β (P=0.0061), GM-CSF (P=0.0265), TNF-α (P=0.0237), and CXCL2 (P=0.0191), and lower levels of IL-17A (P=0.0098) and CXCL9 (P=0.0353) (Figure 3A, C). Similarly, HC monocytes produced higher levels of IL-1β (P=0.0053) and CXCL2 (P=0.0191), and lower levels of IL-17A (P=0.0018) than monocytes from TBI individuals.

Following iMtb stimulation, monocytes from both TBI and TBD groups showed significantly increased production of several cytokines and chemokines compared with HC, including IL-1α (TBI: P=0.0153; TBD: P=0.0228), IL-8 (TBI: P=0.0129; TBD: P=0.0278), IL-23 (TBI: P=0.0158; TBD: P=0.0400), and GM-CSF (TBI: P=0.0346; TBD: P=0.0203) (Figure 3B, D; Supplemental Figure 1). Notably, despite comparable baseline cytokine production, only monocytes from TBI individuals showed significantly increased levels of IL-1RA (P=0.0084), IL-17A (P=0.0171), IL-18 (P=0.0438), IFN-γ (P=0.0382), CXCL1 (P=0.0194), CXCL9 (P=0.0129), and CCL2 (P=0.0420) compared with HC (Figure 3B, E). These findings demonstrate that while both TBI and TBD monocytes respond robustly to iMtb stimulation, monocytes from individuals with TBI exhibit a broader and more distinct cytokine and chemokine response compared with healthy controls. This suggests that TBI monocytes may be functionally distinct from TBD monocytes, enabling a heightened response to iMtb. In contrast, the blunted baseline and narrower inducible responses observed in TBD may reflect monocyte dysfunction or exhaustion associated with active disease.

### Reduced mitochondrial metabolism in monocytes from TBI and TBD individuals

We next compared the metabolic function of monocytes from HC, TBI, and TBD individuals using the SCENITH assay, which measures puromycin incorporation as a proxy for protein synthesis, a process that accounts for approximately 50% of a cell’s total metabolic activity (Figure 4A). The addition of metabolic inhibitors 2-DG and oligomycin enabled us to evaluate cellular dependence on glycolysis and mitochondrial respiration, respectively.

**Figure 4.**
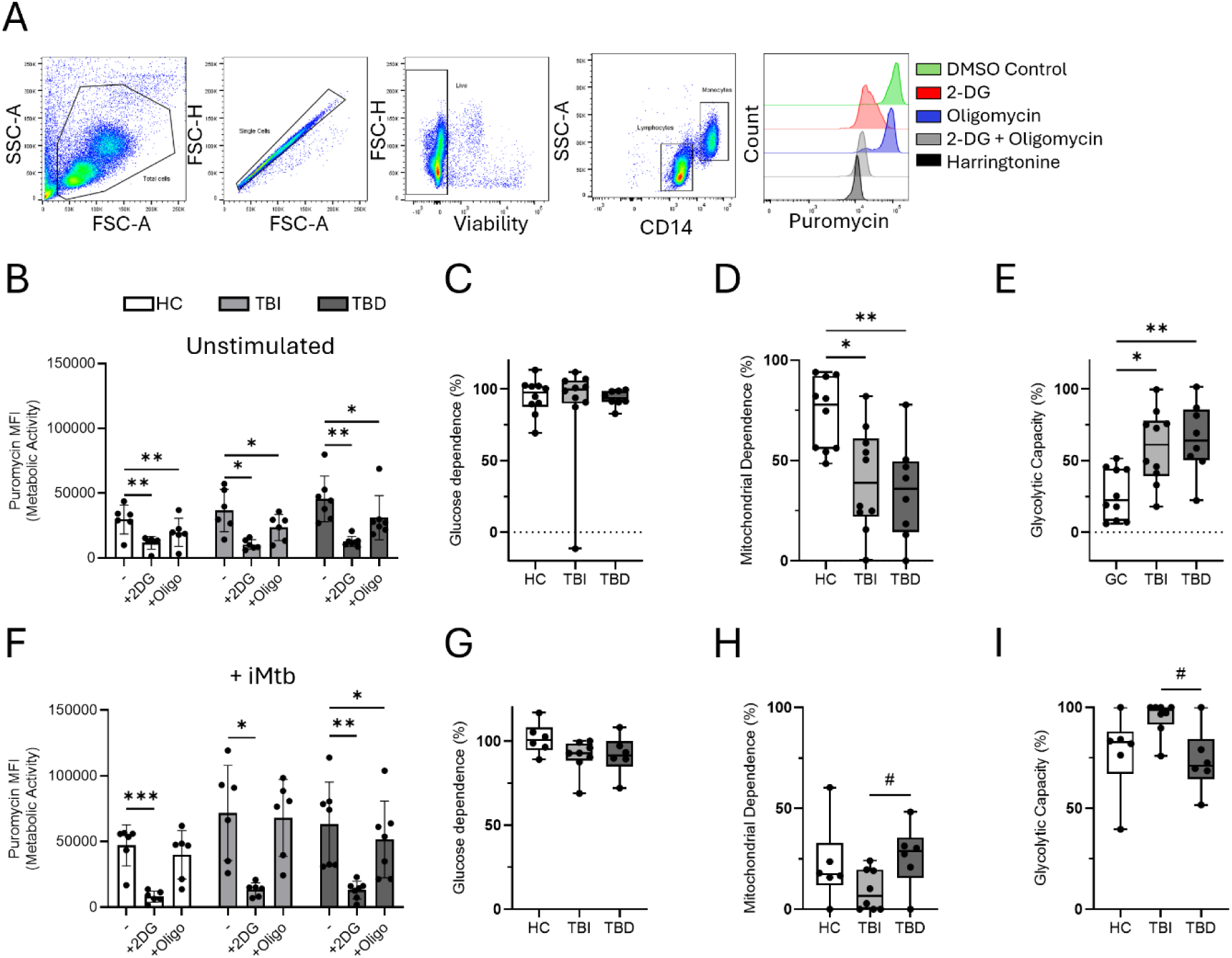
Monocyte metabolic response to Mtb is distinct in TBI. PBMC were isolated from peripheral blood and cultured with or without irradiated H37Rv (iMtb) 10ug/mL for 18 hr. Post-18hr PBMC cultures were treated with DMSO control or metabolic inhibitors 2-deoxyglucose (2DG), oligomycin (Oligo), or combinations of both, or harringtonine, which inhibits protein synthesis and acts as a negative control for this assay. Puromycin was then added, and cells were cultured for 40 min. Cells were washed and stained with fluorochrome-conjugated antibodies against CD14 and puromycin and analyzed by flow cytometry. Puromycin MFI levels measure overall protein synthesis and act as a surrogate for metabolic activity. **A)** Representative flow cytometry gating strategy showing the analyses of monocytes and their puromycin staining across metabolic treatments. **B)** Puromycin MFI values measuring overall metabolic activity in unstimulated monocytes across untreated (-, DMSO control), 2DG, and oligomycin treatments, from HC (n=10) and people with TBI (n=10) and TBD (n=8). **C-E)** Unstimulated monocytes percentage glucose (C) and mitochondrial (D) dependence and glycolytic capacity (E) across all cohorts. **F)** Puromycin MFI values measuring overall metabolic activity in iMtb-stimulated monocytes across untreated, 2DG, and oligomycin treatments from HC (n=6) and people with TBI (n=8) and TBD (n=6). **G-I)** iMtb-stimulated monocytes percentage glucose (G) and mitochondrial (H) dependence and glycolytic capacity (I) across all cohorts. P values generated using Two-Way ANOVA with Dunnett’s (B, F), Kruskal Wallis test with Dunn’s multiple comparison (D, E) and Mann Whitney test (H).

In unstimulated conditions, monocyte metabolic activity (measured by puromycin mean fluorescence intensity [MFI]) was reduced following treatment with either 2-DG or oligomycin across all groups (Figure 4B), confirming sensitivity to both glycolytic and oxidative metabolic pathways.

Glucose dependency, as calculated by SCENITH, was comparably high in monocytes from HC, TBI, and TBD individuals (Figure 4C), indicating a shared reliance on glucose to fuel glycolysis under basal conditions. In contrast, mitochondrial dependency was significantly reduced in monocytes from both TBI (P = 0.01) and TBD (P = 0.0078) individuals compared with HC (Figure 4D), suggesting a metabolic shift away from oxidative phosphorylation. This reduction in mitochondrial dependence is consistent with a compensatory increase in glycolytic capacity (Figure 4E).

### TBI monocytes exhibit enhanced glycolytic response to Mtb

Given the distinct clinical and immunological profiles of TBI and TBD, we hypothesized that monocyte metabolic responses to Mtb would differ between TBI and TBD individuals. Following iMtb stimulation, treatment with 2-DG significantly reduced monocyte metabolic activity in all groups, confirming a strong dependence on glucose to fuel the activation-induced response (Figure 4F–G).

In contrast, oligomycin treatment had no significant impact on metabolic activity post-stimulation (Figure 4F), and calculated mitochondrial dependency remained low across all groups (Figure 4H). Notably, monocytes from TBI individuals exhibited significantly lower mitochondrial dependence and higher glycolytic capacity compared with those from TBD individuals (P = 0.031), indicating a more flexible metabolic profile.

Collectively, these data reveal a clear divergence in metabolic responsiveness to Mtb between TBI and TBD monocytes. Monocytes from TBI individuals display enhanced glycolytic capacity and reduced reliance on mitochondrial metabolism, both at baseline and in response to stimulation; hallmarks of a metabolically reprogrammed, activated phenotype. In contrast, the limited metabolic adaptability observed in monocytes from TBD individuals may reflect impaired functional reprogramming during active disease.

### Treatment restores monocyte mitochondrial function toward a healthy baseline

To determine whether treatment influenced monocyte phenotype or metabolic function, we further stratified participants based on treatment status (Figure 5). Individuals with TBI were grouped as pre-treatment (TBI/-) or on-treatment (TBI/+), and those with TBD were classified into intensive-phase treatment (TBD/I, <2 months, multi-drug) or continuation-phase treatment (TBD/C, >2 months, reduced-drug) cohorts. CD14 expression is significantly elevated in TBI/+ (P=0.0228) and TBD/I (P=0.0069) individuals (Figure 5A). CD45RA expression levels were significantly increased in TBI/+ (P=0.0208), TBD/I (P=0.0310), and TBD/C (P=0.0078; Figure 5B). Of note, HLA-DR expression is only significantly decreased in TBD/I (P=0.0069) and TBD/+ (P=0.0023) individuals (Figure 5D), and TNFRI expression is only reduced in TBI/+ individuals (P=0.0162; Figure 5E).

**Figure 5.**
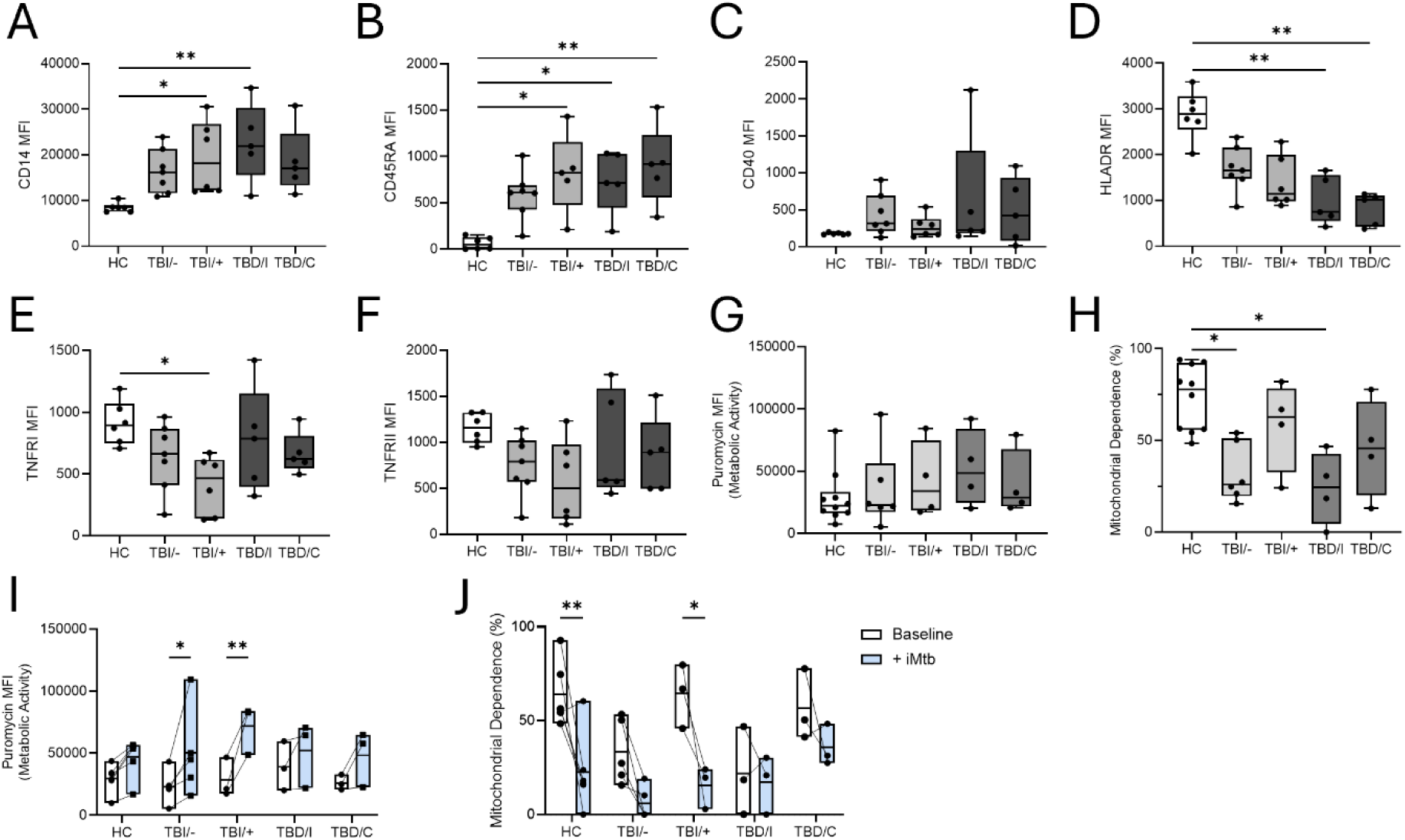
Treatment impact on monocyte phenotype and function. PBMC were isolated from peripheral blood and analysed by flow cytometry or cultured with or without irradiated H37Rv (iMtb) 10ug/mL for 18 hr. **A-F)** Expression levels of CD14 (A), CD45RA (B) and CD40 (C) and HLA-DR (D), TNFRI (E) and TNFRII (F) on classical monocytes ex vivo. Post-18hr PBMC cultures were treated with DMSO control or metabolic inhibitors 2-deoxyglucose (2DG), oligomycin (Oligo), or combinations of both, or harringtonine, which inhibits protein synthesis and acts as a negative control for this assay. Puromycin was then added, and cells were cultured for 40 min. Cells were washed and stained with fluorochrome-conjugated antibodies against CD14 and puromycin and analyzed by flow cytometry. Puromycin MFI levels measure overall protein synthesis and act as a surrogate for metabolic activity. **G)** Puromycin MFI values measuring overall metabolic activity in unstimulated, DMSO control-treated, monocytes in HC (n=10), TBI pre-treatment (TBI/-, n=6) and TBI on-treatment (TBI/+, n=4) and TBD in the intensive (TBD/I, n=4)) and continuation (TBD/C, n=4) treatment phase. **H)** Percent mitochondrial dependence in unstimulated monocytes. **I)** Puromycin MFI values measuring overall change in metabolic activity from unstimulated (white) to iH37Rv-stimulated (blue) monocytes. **J)** Change in percentage mitochondrial dependence in iH37Rv (iMtb)-stimulated monocytes. P values generated using Two-Way ANOVA with Sidak’s multiple comparison test.

Assessment of their metabolic activity revealed no differences in baseline protein synthesis between these subgroups (Figure 5G). However, monocytes from TBI/-individuals showed significantly reduced mitochondrial dependence compared with HC, while those from TBI/+ did not (P = 0.0258; Figure 5H). Similarly, TBD/I monocytes displayed reduced mitochondrial dependence compared with HC (P = 0.0198), whereas mitochondrial dependence in TBD/C monocytes was not significantly different from HC (Figure 5H).

Upon iMtb stimulation, only monocytes from TBI individuals significantly increased their overall metabolic activity (TBI/−: P = 0.0137; TBI/+: P = 0.0024; Figure 5I). Again, a corresponding decrease in mitochondrial dependence was only observed in TBI individuals on treatment (P=0.0185) and HC (P=0.0044; Figure 5J). In contrast, monocytes from TBD individuals showed no significant change in either metric following iMtb stimulation, regardless of treatment phase (Figure 5I-J).

These data suggest that both TBI and TBD are associated with reduced mitochondrial metabolism in circulating monocytes, particularly in untreated individuals, and that both TB preventative treatment (TPT) and TBD treatment may partially restore metabolic function toward a healthy baseline.

## Discussion

Our study reveals that Mtb infection induces distinct alterations in circulating monocyte phenotype, function, and metabolism, that differ between individuals with TBI and TBD. Despite similar monocyte subset frequencies across groups, phenotypic profiling showed that monocytes in TBI individuals exhibit an upregulation of activation markers (e.g., CD14, CD45RA) alongside downregulation of HLA-DR and TNF receptors, key mediators of antigen presentation and inflammation. Notably, HLA-DR expression declined progressively from TBI to TBD, suggesting a trajectory of monocyte dysfunction with disease progression which has been highlighted as a biomarker of infection in TB, sepsis and paediatric infection (24–28). Functionally, monocytes from both groups remained responsive to iMtb stimulation, but TBI monocytes exhibited a more robust cytokine and metabolic response compared with TBD monocytes.

Among the analytes elevated in TBI compared with TBD and healthy controls, IL-1RA, IL-17A, and CXCL9 showed the most pronounced increase, unaffected by outlier values. IL-1RA, an antagonist of IL-1α/β signalling produced predominantly by monocytes and neutrophils, has been proposed as a biomarker for TB spectrum stratification (29). In our study, IL-1RA was significantly upregulated in monocytes from individuals with TBI, but not from those with TBD, supporting a potential role for monocyte-derived IL-1RA in maintaining infection containment. However, previous findings on IL-1RA expression in TB have been inconsistent, with some studies reporting higher IL-1RA levels in response to TB antigens in TBD relative to TBI, and others showing the opposite (30–33). Notably, these prior studies were based on whole blood or PBMC cultures, making it difficult to resolve cell-specific contributions. By focusing on purified monocytes, our data provide more direct evidence that monocyte-specific IL-1RA production may differentiate TBI from TBD.

Similarly, we observed significant upregulation of IL-17A and CXCL9 in monocytes from TBI but not TBD individuals (Figure 1E). CXCL9 acts as a critical chemoattractant for CXCR3⁺ T cells, facilitating the recruitment and activation of IFN-γ–producing effector T cells (34). IL-17A is a potent inducer of chemokines that recruit immune cells such as T cells and neutrophils, and stimulates antimicrobial peptide production from epithelial cells (35). Collectively, this monocyte cytokine and chemokine profile characterized by elevated IL-1RA, IL-17A, and CXCL9 in TBI may reflect an immunological environment conducive to effective containment of Mtb infection.

We hypothesize that the distinct effector cytokine and chemokine profile observed in monocytes from individuals with TBI is underpinned by metabolic reprogramming that favours glycolysis over mitochondrial respiration. TBI monocytes displayed enhanced glycolytic capacity and reduced mitochondrial dependence both at rest and upon stimulation, consistent with an activated immunometabolic phenotype (Figure 4D-E). In contrast, TBD monocytes failed to undergo this shift, suggesting impaired metabolic plasticity which supports previous reports of their overall diminished bioenergetics (20). Given the known link between glycolysis and effector function, this metabolic rewiring is likely associated with the enhanced production of cytokines and chemokine observed specifically in TBI monocytes, which may facilitate the effective immune containment of Mtb. Of note, to assess differential sustained metabolic responses of monocytes to Mtb between TBI and TBD individuals, cells were cultured for 18 hours with iMtb followed by the SCENITH protocol. We acknowledge that future studies incorporating Seahorse Analyzer assays on freshly isolated monocytes may provide further insight into the dynamic and immediate metabolic changes that occur upon iMtb stimulation.

These data resonate with and extend the work of other research groups investigating host metabolic responses in TB. Notably, bioenergetic dysfunction of monocytes in South African TBD cohorts have been reported (20). In addition, myeloid cell metabolic rewiring governs innate immune cell function in the context of BCG vaccination (36) and heterologous innate immunity to Mtb elicited by adenoviral vectored COVID-19 vaccination (37). Our findings complement and build upon these studies by focusing specifically on purified circulating monocytes metabolic capacity and effector cytokine output across the TB spectrum.

Importantly, our findings also indicate that treatment, including TPT in TBI and multidrug regimens in TBD, partially restores mitochondrial function in circulating monocytes, shifting their metabolic baseline profile closer to that of healthy individuals. This suggests that monocyte metabolism is dynamic and responsive to therapeutic intervention, indicating that mitochondrial dependency metrics hold potential as a novel biomarker for treatment efficacy and disease resolution. Moreover, these insights underscore the therapeutic potential of targeting monocyte metabolic pathways to enhance host immune function in TB.

These results contribute to a growing body of evidence that immunometabolism shapes infection outcomes in TB. Since circulating monocytes give rise to macrophages, including lung-resident alveolar macrophages, targeting metabolic pathways in monocytes may enhance both peripheral and pulmonary immune responses (38–40). Although our study focused on peripheral blood monocytes, it is reasonable to hypothesize that some aspects of monocyte immunometabolic reprogramming are reflected in lung-resident or recruited mononuclear phagocytes, given the dynamic trafficking of monocytes from blood to tissue. However, the lung microenvironment may also induce additional, tissue-specific metabolic adaptations that could modify these peripheral phenotypes. Recent evidence indicates that human airway macrophages are metabolically and functionally distinct from monocyte-derived macrophages in response to Mtb (41,42) and murine studies highlight the metabolic heterogeneity across alveolar and interstitial lung macrophages in response to Mtb (18). If future studies reveal differences between systemic and local immune programming, this will underscore compartmentalized immune responses, helping to delineate distinct mechanisms of immune protection or pathogenesis in TB. Importantly, such insights reinforce the potential of monocyte metabolism as a promising target for host-directed therapies, particularly those aimed at preventing progression from TBI to TBD by enhancing both systemic and lung-localized immunity.

While our study provides novel insights into monocyte immunometabolism across the TB spectrum, several limitations should be acknowledged. First, the relatively small sample size may limit the generalizability of our findings and the statistical power to detect more subtle differences or correlations with clinical outcomes. Second, we did not establish a direct link between the observed monocyte metabolic and phenotypic changes and disease progression or treatment success, which will require longitudinal studies with larger cohorts. Third, our functional assays utilized irradiated H37Rv. This well-characterized lab strain facilitates standardization however, future studies incorporating live, virulent strains are essential to fully elucidate how pathogen variability shapes host monocyte responses. Despite these limitations, our data provide a valuable foundation for future investigations aimed at elucidating the functional benefits of this metabolic plasticity in TBI, and its impact on bacterial killing. Such studies will be essential for identifying robust biomarkers and therapeutic targets through detailed mechanistic analyses of monocyte immunometabolism in tuberculosis.

In summary, our data reveal that Mtb infection drives distinct immunometabolic changes in circulating monocytes with TBI characterised by functional responsiveness and glycolytic, pro-inflammatory profile, while TBD is marked by impaired metabolic plasticity. Furthermore, we show that TB treatment of both TBI and TBD individuals may restore monocyte mitochondrial function. These findings offer important insight into the cellular mechanisms of immune containment in TB and support the rationale for treating TBI. Moreover, they underscore the potential of monocyte metabolism as a biomarker of disease state and a tractable target for host-directed therapies. Continued research building on these insights will be crucial for advancing TB diagnosis, monitoring treatment efficacy, and developing novel interventions to improve patient outcomes.

## Acknowledgements

We sincerely thank all the individuals who participated in this study. We are especially grateful to the patients for their time, generosity, and willingness to contribute to research aimed at improving the understanding and treatment of tuberculosis. Their participation made this work possible. The following reagent was obtained through BEI Resources, NIAID, NIH: Mycobacterium tuberculosis, Strain H37Rv, Gamma-Irradiated Whole Cells, NR-49098. We gratefully acknowledge the support of the Core Flow Cytometry Facility and Prof. Nollaig Bourke’s research team at the Trinity Translational Medicine Institute.

---

**Supplemental Figure 1.**
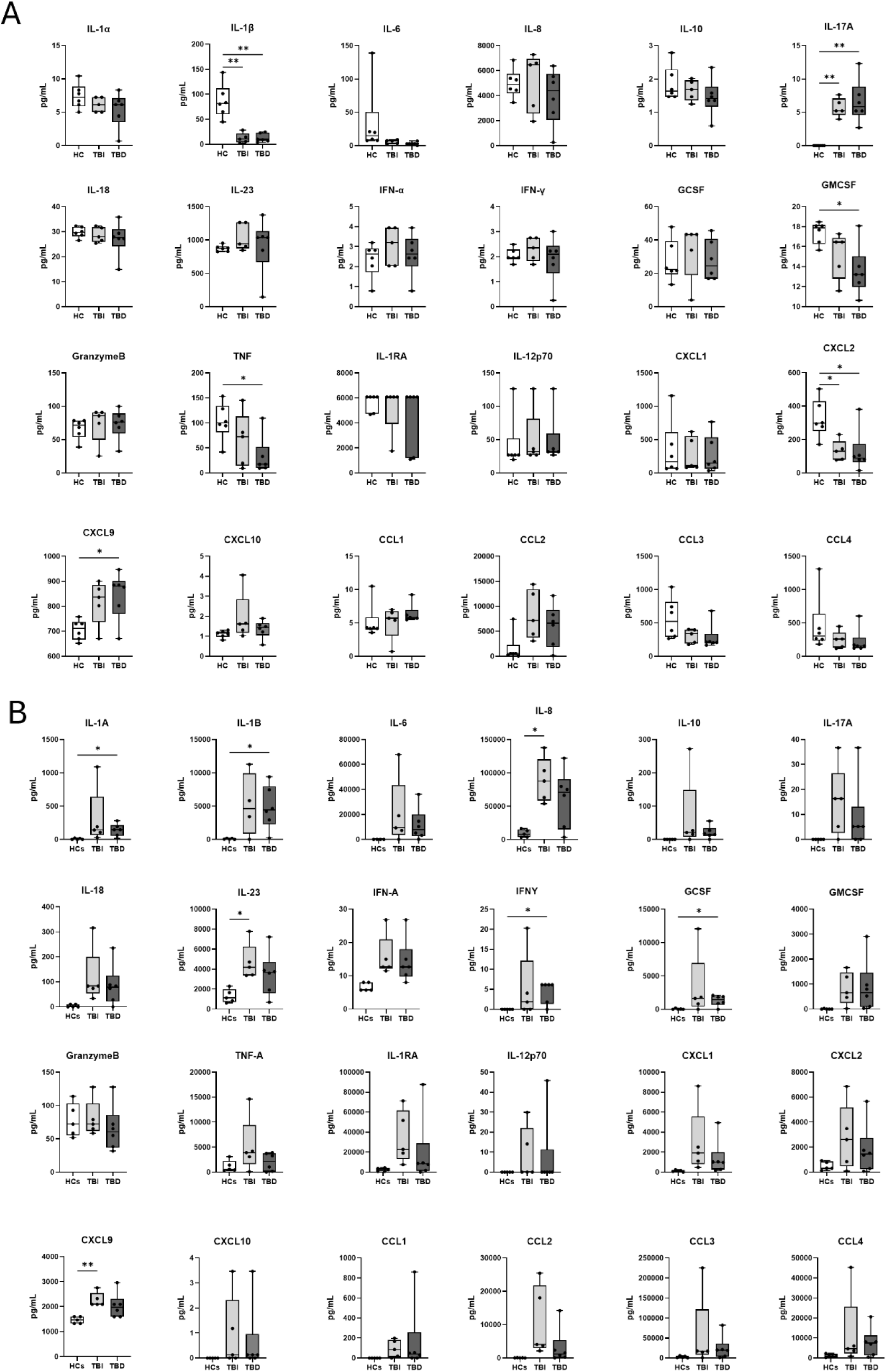
Raw concentration values for monocyte cytokine and chemokine production. Monocytes were isolated from PBMC and cultured for 18 hrs with and without irradiated H37Rv (iMtb). **A-B)** Concentration of cytokines and chemokines (pg/mL) in supernatants collected from unstimulated (A) and iMtb-stimulated (B) monocyte cultures. P values generated using 2 Way ANOVA with Tukey’s multiple comparison test.

**Supplemental Figure 2.**
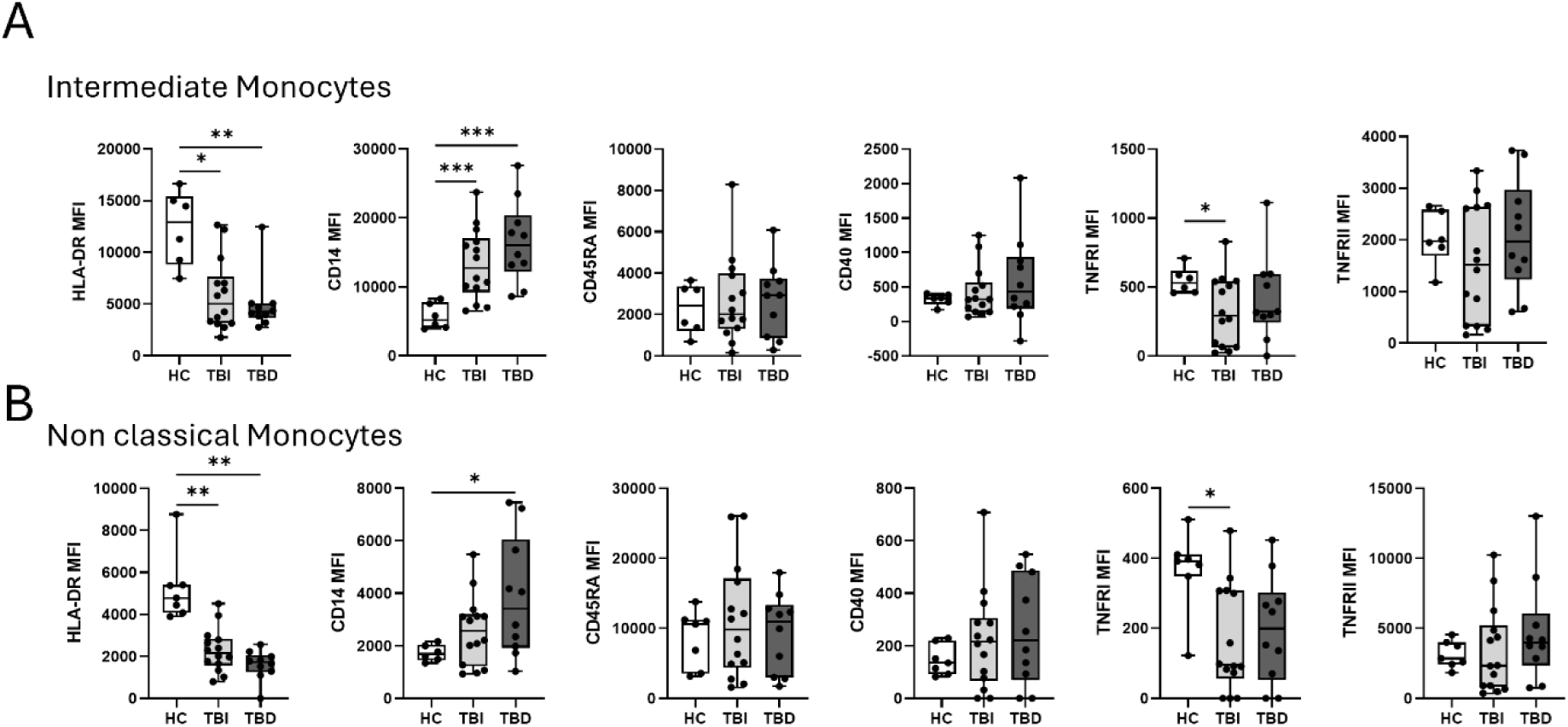
TB infection alters intermediate and non-classical monocyte phenotype. PBMC were isolated from peripheral blood and stained for flow cytometry analysis. **A-B)** MFI values for CD14, HLA-DR, CD45RA, CD40, TNFRI, and TNFRII expression on intermediate (A) and non-classical (B) monocytes across HC, TBI and TBD. P values generated using Two-Way ANOVA with Tukey’s multiple comparison test.

## Notes

**Sources of Support:** This work was funded by the Health Research Board Ireland (HRB-ILP-POR-2022-033 and HRB-EIA-2019-010; awarded to S.A.B.) and The Royal City of Dublin Hospital Trust (RCDH app 185; Awarded to J.K.). The following reagent was obtained through BEI Resources, NIAID, NIH: Mycobacterium tuberculosis, Strain H37Rv, Gamma-Irradiated Whole Cells, NR-49098).

### Competing Interest Statement

The authors have declared no competing interest.

### Summary of Updates

Added discussion points and alterations to how data was presented.

